# Ongoing loss of viable neurons for weeks after mild perinatal hypoxia-ischemia

**DOI:** 10.1101/2024.12.19.629457

**Authors:** Melanie A. McNally, Lauren A. Lau, Simon Granak, David Hike, Xiaochen Liu, Xin Yu, Rachel A. Donahue, Lori B. Chibnik, John V. Ortiz, Alicia Che, Frances Northington, Kevin Staley

## Abstract

Mild hypoxic-ischemic encephalopathy is common in neonates with no evidence-based therapies, and 30-40% of patients experience adverse outcomes. The nature and progression of mild injury is poorly understood. Thus, we studied the evolution of mild perinatal brain injury using longitudinal two-photon imaging of transgenic fluorescent proteins as a novel readout of neuronal viability and activity at cellular resolution. In vitro, perinatal murine organotypic hippocampal cultures underwent 15-20 minutes of oxygen-glucose deprivation. In vivo, mild hypoxia-ischemia was completed in post-natal day 10 mouse pups of both sexes with carotid ligation and 15 minutes of hypoxia. Consistent with a mild injury, minimal immediate neuronal death was seen and there was no volumetric evidence of injury by ex vivo MRI 2.5 weeks after injury. In both the hippocampus and neocortex, these mild injuries resulted in a significantly delayed and progressive neuronal loss in the second week after injury, measured by fluorophore quenching. Mild hypoxia-ischemia transiently suppressed cortical network activity followed by normal maturation. No post-injury seizures were seen. The participation in network activity of individual neurons destined to die was indistinguishable from those that survived for 4 days post-injury. In conclusion, our results showed that mild perinatal brain injury resulted in a prolonged increase of neuronal death. Neurons that died late were functioning normally for days after injury, suggesting a new pathophysiology of neuronal death. Critically, the neurons destined to die late demonstrated multiple biomarkers of viability long after mild injury, suggesting their later death may be modified with neuroprotective interventions.

**SIGNIFICANCE STATEMENT:** Neonatal encephalopathy due to peripartum hypoxia-ischemia (HI) is a major cause of neonatal mortality and morbidity worldwide. Of these infants, most are categorized as having mild HI. Infants with mild HI have significant long-term disabilities. There are currently no evidence-based therapies, largely because the progression and pathophysiology of mild injury is poorly understood. We have identified, for the first time, that mild perinatal HI results in a delayed and prolonged increase in neuronal death. The cortical and hippocampal neurons that die over a week after injury participate normally in neural network activity and exhibit robust viability for many days after injury, indicating a novel pathophysiology of neuronal death. Clinically, these data suggest an extended therapeutic window for mild perinatal HI.

## INTRODUCTION

Neonatal encephalopathy due to intrapartum hypoxia-ischemia is a major cause of neonatal mortality and morbidity with 2-8 in 1000 live births being affected worldwide (Kurinczuk et al., 2010). Of these infants, most are categorized as having mild encephalopathy (Lee et al., 2013). Early studies found that term neonates with mild hypoxic-ischemic encephalopathy (HIE) had similar long term neurodevelopmental outcomes to neonates without HIE (Sarnat and Sarnat, 1976; Robertson et al., 1989). However, mild HIE is now diagnosed within the first few hours after birth to determine eligibility for therapeutic hypothermia, which is reserved for moderate to severe encephalopathy (Azzopardi et al., 2009). Infants with mild HIE have significant disabilities compared to unaffected peers, with up to 25% having poor outcomes by 18 months and 35-40% by 5 years of age (Van Kooij et al., 2010; Chalak et al., 2018; Conway et al., 2018). Thus, the long-term global burden of disability for these patients is significant and there are currently no therapeutic interventions with demonstrated benefit. To develop therapies, we first need to understand the progression of injury in these patients (Sarnat et al., 2020; McDouall et al., 2022).

Injury progression after *moderate to severe* perinatal hypoxia-ischemia (HI) has been studied with various modalities in preclinical models and in patients, all having limitations (Nakajima et al., 2000; Geddes et al., 2001; Northington et al., 2011b). The relationships between the number of visible dying neurons, the rate of neuronal death, the rate of efferocytosis (i.e. clearance) of dying neurons, and the time elapsed since injury are complex (Cevik and Dalkara, 2003; Northington and Martin, 2012; Zille et al., 2012; Balena et al., 2023). Histopathological assays for neuronal death are limited in their ability to achieve the temporal resolution required to quantify this injury and test for progression in real time. Longitudinal studies of neuronal death in living tissue are needed for accurate quantification (Galluzzi et al., 2015; Balena and Staley, 2024). Progression of injury is a particularly important question in mild HIE, where the possibility of late neuronal death might alter acute treatment decisions.

Injury progression after mild perinatal HI has not yet been elucidated in preclinical studies. Clinically, MRI is used, but it is not sufficiently sensitive to assess for the degree and progression of injury in these patients (Conway et al., 2018). The acute MRI abnormalities detected in ∼20-40% of contemporary mild HIE cohorts are subtle, most often with only small areas of signal change in the white matter or cortex (Prempunpong et al., 2018; Walsh and Inder, 2018). The association of more subtle injury seen on MRI in infants with mild HIE to childhood outcomes is unknown. EEG has also been used to assess HIE (Gluckman, 2005; Murray et al., 2009). While its utility studying disease progression after mild HIE is unknown, EEG shows acute abnormalities in up to 70% of infants with mild HIE (Garvey et al., 2021). These findings indicate acute dysfunction of injured neural networks. At a cellular level, injured neurons are subject to synaptic stripping (Cullheim and Thams, 2007; Kettenmann et al., 2013) and loss of postsynaptic spines (Waataja et al., 2008; Guo et al., 2012), which functionally disconnect the injured neurons from the neural network. The degree of participation of individual neurons in network activity should therefore be a sensitive measure of viability when measuring injury progression longitudinally.

In this study, we used longitudinal assays in vitro and in vivo to examine sequelae of perinatal injury. Viability of individually tracked neurons was assessed using ongoing expression of transgenic fluorescent proteins (Arrasate et al., 2004; Linsley et al., 2019; Balena et al., 2023), and participation in network activity (Lau et al., 2022). With these tools, we demonstrated a very delayed and progressive neuronal loss following mild perinatal HI that occurred following a preceding period of normal neuronal function, potentially opening a therapeutic window for interventions for mild HIE to prevent life-long adverse outcomes.

## MATERIALS AND METHODS

### Animals

All animal protocols were approved by the Massachusetts General Hospital Institutional Animal Care and Use Committee (protocol #2020N000141). All studies were conducted in accordance with the United States Public Health Service’s Policy on Humane Care and Use of Laboratory Animals. Wild type mice (C57bl/6; Jackson Labs) of either sex were used. On post-natal day 1 (P1), all animals underwent intracerebroventricular injection in the left hemisphere with pAAV1-hSyn1-mRuby2-GSG-P2A-GCaMP6s-WPRE-pA (Addgene, #50942). Mouse pups remained in the home cage with the dam under standard husbandry conditions until P6–8 when organotypic hippocampal slice cultures were prepared. For in vivo experiments, pups stayed with their dams under standard husbandry conditions until P7-8 when the cranial window was placed. They were removed from their home cages for subsequent imaging and interventions, but returned to their dam between timepoints through P28 at the experimental endpoint.

### In vitro organotypic slice cultures, oxygen-glucos deprivation, and imaging

Organotypic hippocampal slice cultures were prepared as membrane insert-mounted cultures (Stoppini et al., 1991). Briefly, hippocampi were obtained from P6-8 mice, cut to 400mm slices, transferred to a 6-well dish containing a membrane insert (PICMORG50; Millipore-Sigma), fed twice weekly with neurobasal-A media supplemented with 500mM Glutamax, 2% B-27 and 0.03mg/mL gentamycin (Invitrogen), and incubated at 35°C in a 5% CO_2_ incubator. Cultures were imaged starting at 3-4 days in vitro (DIV3-4) within a TC-MIS miniature incubator connected to a TC-1-100-I temperature controller set to the same atmospheric conditions (Bioscience tools) and randomly assigned to experimental groups. For oxygen-glucose deprivation (OGD) at DIV3-4, the media was switched to one without glucose and the chamber was aerated with 95% nitrogen/5% CO_2_ for 15-20 minutes after a 5-minute equilibration period. Longer durations of OGD result in pronounced cell damage and inability to image the slice culture chronically to study injury progression, whereas shorter durations do not produce evidence of injury by caspase activation, neuronal swelling, or cell death (Bahari et al., 2024). For caspase experiments, slices were pre-incubated with BioTracker NuvView® Blue Caspase-3 Dye for >30 minutes (Sigma Aldrich, 5μL dye:1 mL media). Two-photon (2P) imaging was performed using a custom-built scanning microscope. 2P images were acquired using custom-designed software (LabVIEW), a scan head from Radiance 2000 MP (Bio-Rad) equipped with a 40X, 0.8NA water-immersion objective (Olympus), and a mode-locked Ti:Sapphire laser (MaiTai; Spectra-Physics). The objective was mounted to an inverter arm positioned beneath the miniature incubator. During imaging, mRuby was excited at 760nm, green fluorescent protein (GFP) was excited at 910 nm, and blue caspase dye fluorescence was excited at 820 nm. Emission was detected through three filters: 470/50, 545/30, and 620/100 nm. Images were reconstructed offline in ImageJ (RRID: SCR-003070).

### In vivo murine cortical window placement

Using a modified method based on that of Che et al. (2021), cortical windows were placed in P7-8 mouse pups under sterile conditions. Briefly, isoflurane anaesthetized mice were placed on a warmer. The scalp was washed with alternating solutions of 70% ethanol and 10% povidone-iodine. A large section of scalp was removed between the ears to expose both hemispheres and the skull was dried. A custom-made, titanium head plate with an inner diameter of 5 mm was centered over the right somatosensory cortex and adhered with veterinary adhesive (Metabond®). A craniotomy was performed with gentle etching using a 0.75mm lancet (World Precision Instruments, #504072), bleeding was controlled using epinephrine hemostatic pellets (Pascal). A 3mm cranial window (Thomas Scientific, #1217N66) was placed over the exposed cortex and subsequently fixed to the skull first with Vetbond®, and then with a second layer of Metabond®. Total anesthesia duration was <60 minutes per pup. Intra-op and post-op analgesia with carprofen (Zoetis) was administered.

### Neonatal hypoxic-ischemic brain injury

In P10 mice, a modified Vannucci model was used to induce HI (Burnsed et al., 2015; McNally et al., 2019). P10 was chosen because it most closely mimics human term neonate brain maturation (Semple et al., 2013). Under <10 minutes of isoflurane anesthesia, unilateral ligation of the right carotid artery was completed followed by a 1-hour recovery period. Pups were subsequently exposed to 15 minutes of hypoxia at 36°C (FiO_2_, 0.08). 15 minutes was the longest duration of hypoxia that reliably permitted chronic imaging of the cortex under the cranial window. Higher rates of mortality and widespread acute cell death manifest by nearly complete loss of neuronal fluorescence emission were a frequent complication after longer periods of hypoxia with this experimental design. Control sham pups underwent the same isoflurane anesthesia exposure, and the right carotid artery was exposed, but not ligated. They underwent the same recovery period and time away from dam. Pups were randomly assigned to experimental groups. In the hands of an experienced practitioner (>2 years), 80-85% survival was achieved through the experimental end point (P28) with an overall 65% success rate (some cranial windows were not of sufficient quality to permit longitudinal imaging). Up to 15% less growth is typical in HI animals compared to sham animals prior to weaning (Burnsed et al., 2015; Diaz et al., 2017; McNally et al., 2019). An exception was one sham animal showed marked weight loss prior to the end of the experiment for unknown reasons and had to be euthanized.

### In vivo murine chronic two-photon imaging

Unanesthetized mouse pups were stabilized by attaching the headplate to a fixed fork beneath the objective. During P10-12 recordings, the pups were placed on a warming pad (rectal temps maintained at 36-37°C). At >P14, pups were placed on a rotary treadmill. A custom-made gantry-type 2P microscope equipped with a MaiTai 80MHz Ti:Sapphire laser (Spectraphysics) and a 16x, 0.8 N.A. water-immersion objective (Nikon) was driven with customized ScanImage software (Vidrio RMR) and used for imaging (Pologruto et al., 2003; Costine-Bartell et al., 2023). High resolution mRuby z-stacks (10 *μ*m step length) of cortical layers 1-3 were acquired of the somatosensory cortex at each timepoint using excitation at 750 nm. 1.1Hz calcium imaging was performed with excitation at 920 nm for 500 frames (7.6 minutes/timepoint). Emission was detected through 605/70 and 525/50 nm bandpass filters (Thorlabs) for mRuby and GCaMP6s, respectively, before PMT detection (Hamamatsu C6438-01).

### Processing and analysis of calcium imaging

Inclusion criteria for calcium imaging included the following: 1. Imaging quality sufficient to resolve individual neuronal activity in layer II/III somatosensory cortex and 2. minimum of 24h of data. Animals were excluded from all analyses if atypical injury was noted (eg. no imageable tissue under cortical window at P11 or later timepoints, or no neuronal or network activity with sustained high basal calcium signal at P11 or later timepoints). Calcium imaging was processed and analyzed using modified methods based on Lau et al. (2022) and Che et al. (2021). Timelapse recordings were first motion corrected using the Image Stabilizer plugin in Fiji. Epochs where movement was too significant to correct were removed. For network analyses, The Fiji plugin TrackMate(Schindelin et al., 2012) was used to automatically select soma ROIs of active, GCAMP+ cells from the frame-to-frame standard deviation (STD) projection, and calcium signal was extracted as the mean pixel intensity of each ROI. For neuronal analyses of manually tracked neurons, the presence and location of individual neurons were confirmed using aligned mRuby images and manual ROIs were drawn in that location on the STD projection of the calcium imaging (thus, able to capture calcium activity of neurons regardless of activity status). Calcium traces were normalized on a frame-by-frame basis as ΔF/F as: F_raw_ – F_min_ / F_min_. F minimum (F_min_) for each ROI was calculated as the bottom 1^st^ percentile of the mean intensity over all frames. The threshold for being considered in the active state was determined using the kernel density estimation method (Mitani and Komiyama, 2018), with the threshold set at the 90^th^ percentile area under the 1^st^ distribution curve (where the first distribution will largely be comprised of baseline, inactive calcium fluorescence values). To determine what percentage of simultaneously active cells represented a statistically significant network or cortical event, surrogate datasets were constructed by reshuffling activation timing in each cell individually and then averaging the surrogate datasets to create a distribution of event sizes. This was repeated 1000 times and the threshold significance for network or cortical events was then set at *p* < 0.01. The time points where the percentage of cells active exceeded or dropped under the threshold were set to be, respectively, the onset and offset of the network or cortical events. These time points were used to quantify the duration of the events. The maximum percentage of cells simultaneously active during any given network or cortical event was averaged across all network events in each recording to measure ‘network/cortical activation’. Using the times of the network events defined by these analyses, the proportion of network events during which individual neurons were active was measured to define ‘neuronal network participation’.

Of note, neuropil subtraction was completed on a subset of data and no effect on the conclusions of the results for network, cortical, or individual neuronal analyses were seen. Thus, no neuropil subtraction was applied to the shown analyses. Additionally, no video recording was completed during these experiments and thus the state of the animal was unable to be synchronously verified with the recordings.

### Fluorophore quenching assay

A representative subset of neurons was selected on baseline imaging for survival tracking. Inclusion criteria for neuronal survival tracking in vitro included the following: 1. Imaging quality sufficient to resolve individual mRuby-positive neuronal soma in the pyramidal layer of CA1 on experimental day 0 (DIV3/4) at approximately 50 μm below slice surface and 2. same region well imaged for at least 10 days. Inclusion criteria for neuronal survival tracking in vivo included the following: 1. Imaging quality sufficient to resolve individual mRuby-positive neuronal soma in layer II/III somatosensory cortex on P10 and 2. same region well imaged through at least P20. Animals were excluded from all analyses if atypical injury was noted (eg. no imageable tissue under cortical window at P11 or later, or no neuronal or network activity with sustained high basal calcium signal at P11 or later) or if there was spontaneous mortality prior to experimental end point. Every neuron was tracked separately and manually. In vitro and in vivo, the three-dimensional z-stacks from day 0 (P10) through day 10/11 (P20/21) were aligned for each neuron to account for any XYZ drift and verified using blood vessels and surrounding neurons as fiduciary markers. See Figure 2 and Extended Data Figure 2-1 for representative examples. A subset of data was verified using a blinded assessor.

**Figure 1.**
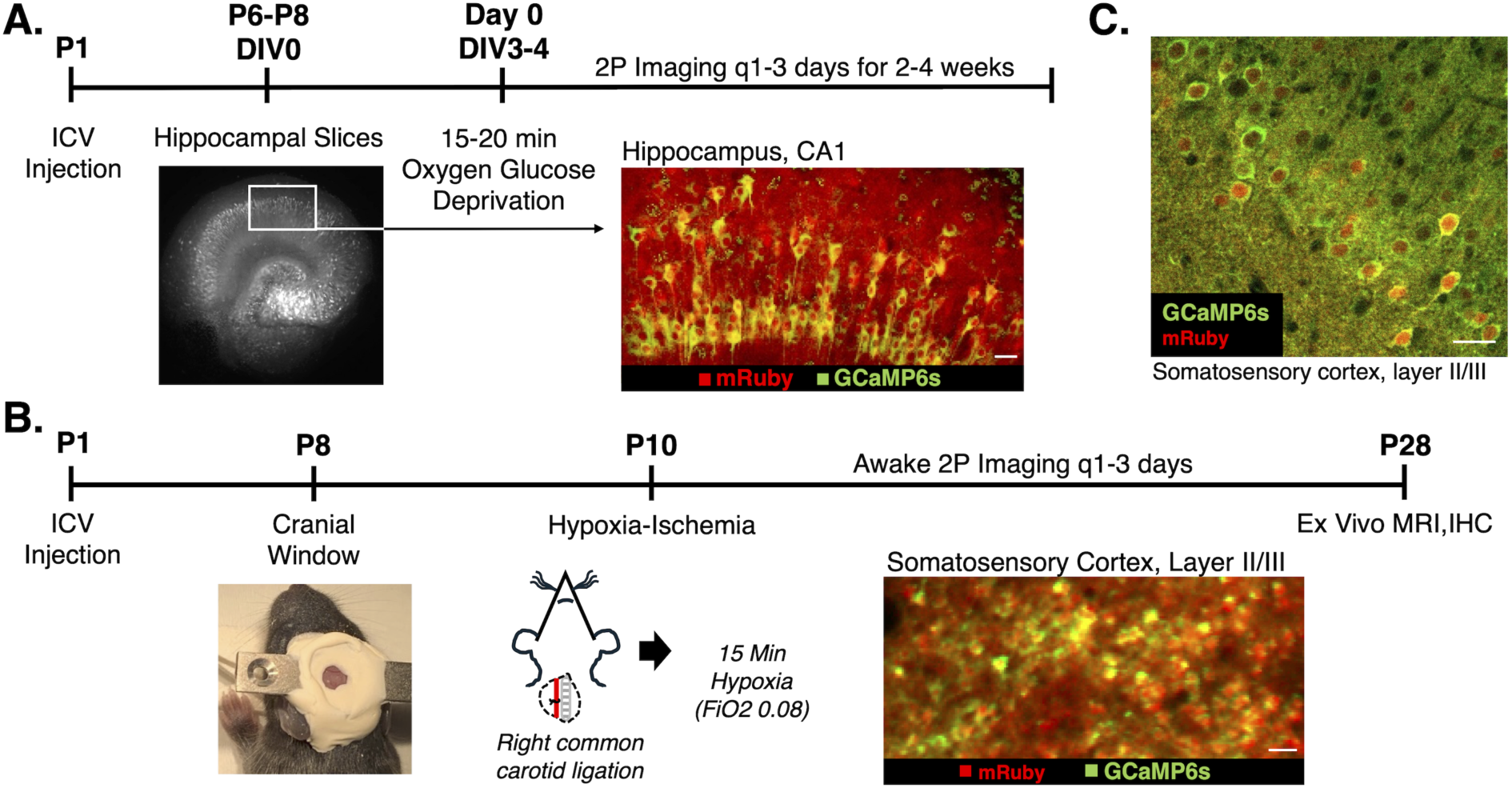
In vivo and in vitro experimental designs. (A) In vitro experimental design with representative hippocampal 2P fluorescence images of *syn*-GCaMP6s expression (standard deviation projection of GCaMP6s activity; white and green) and *syn*-mRuby (red). Scale bar on merge image = 25 μm. (B) In vivo experimental design with image of mouse pup with implanted custom designed titanium head bar and cranial window. Additional representative layer II/III somatosensory cortex 2P fluorescence image of merged *syn*-GCaMP6s standard deviation projection (green) and *syn*-mRuby (red) is shown. Scale bar = 25 μm. (C) Representative immunostaining confocal image of *syn*-driven neuronal GCaMP6s (detected by anti-GFP) and mRuby co-expression in layer II/III somatosensory cortex of a P25 animal post-HI. Manual quantification verified 74% co-expression (*n* = 136 neurons). Scale bar = 25 μm. (C)

**Figure 2.**
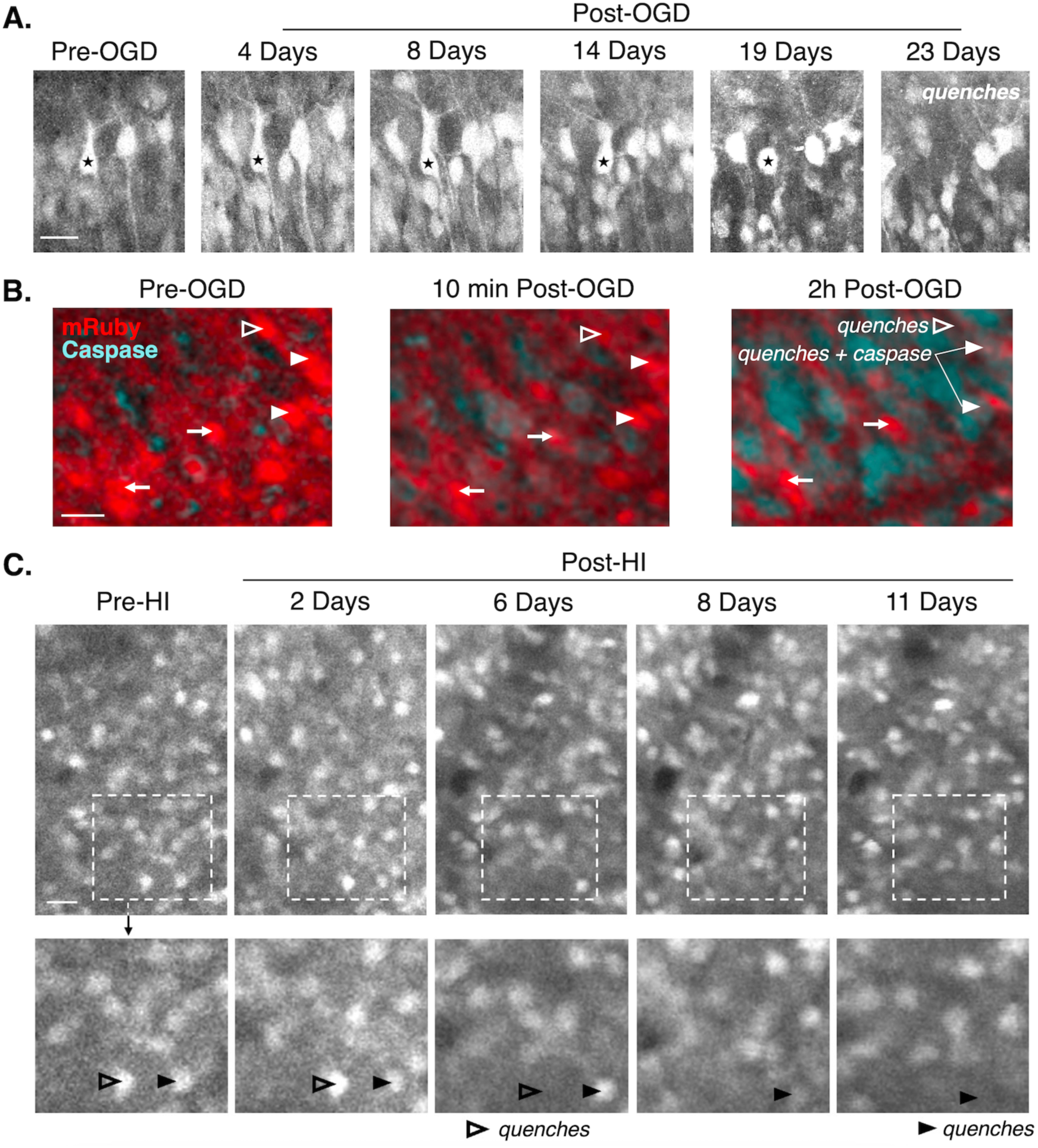
Longitudinal tracking of hippocampal and cortical neuron survival. (A) 2P fluorescence images tracking mRuby-positive hippocampal pyramidal neurons in vitro. The starred neuron showed dendritic retraction and swelling prior to quenching at 23 days. (B) Representative 2P fluorescence images of mRuby-positive hippocampal pyramidal neurons in vitro incubated with NucView® Blue Caspase-3 dye. Neurons that survived post-OGD (arrows) did not show caspase activation acutely. Neurons are shown that quenched 2 hours post-OGD with (filled arrowhead) and without (open arrowhead) caspase activation. (C) Unanesthetized, in vivo 2P fluorescence images of layer II/III somatosensory cortex over time demonstrating tracking of mRuby-positive cortical neurons. Shown are examples of neurons that quenched 6 days post-HI (open arrowhead) and 11 days post-HI (closed arrowhead). Scale bars = 25 μm.

### Magnetic resonance imaging

At P27-29, experimental mice were deeply anesthetized with inhaled isoflurane in preparation for intracardiac perfusion. They were subsequently euthanized by exsanguination with 0.1M phosphate buffered solution (PBS), and then perfused with 4% paraformaldehyde (PFA in 0.1M PBS). Brains were removed and postfixed in the same solution. 1-3 days prior to imaging, brains were washed in PBS alone. The day of imaging, the brains were transferred to Fluorinert^TM^ FC-40 (Sigma Aldrich) and immobilized in plastic centrifuge tubes. Ex vivo ^1^H MRI of the brains was performed on a horizontal 14 tesla (14T) MR scanner (Magnex Sci, UK) located at the Athinoula A. Martinos Center for Biomedical Imaging in Boston, MA. The scanner was equipped with a 14T magnet with a Bruker Avance Neo Console (Bruker-Biospin, Billerica, MA). A 60-mm diameter microimaging gradient insert (Resonance Research, Inc., Billerica, MA) was used providing a peak gradient strength of 1.2T/m. The ex vivo brains were imaged using a Bruker ^1^H quadrature volume coil (ID=35mm). Multi-slice 2D fast spin echo T2-weighted images were acquired utilizing a Rapid Acquisition with Relaxation Enhancement (RARE) sequence with the following parameters: echo time (TE)/repetition time (TR) = 40/6000 ms, RARE-factor = 1, bandwidth = 50000Hz, in-plane resolution of 50µm x 50µm with 400µm slice thickness, 32 slices, and 8 averages for an acquisition time of 3 hours 12 mins. Field of view was 12mm by 12mm with a 240x240 matrix. During imaging, the field of view was adjusted to ensure similar orientation of different samples. The slice package was positioned to cover the entire brain and oriented so the ventral line of the brain was parallel to the ventral side of the slice package.

Data was processed using Analysis of Functional Neuroimages (AFNI). Bruker 2dseq images of the RARE images were converted to AFNI format using ‘to3d’. Data was then resampled using “3dresample” to match the resolution of the Allan Brain Atlas. The Allan Brain Atlas was registered to each data to obtain the ROIs using “3dAllineate”. ROIs based on the Allan Brain Atlas were used to calculate the volume of interested areas.

### Immunohistochemistry

The brain samples were processed by routine histological procedures and cut into 100-μm coronal slices. The anti-GFP antibody (chicken monoclonal, 1:2000, Abcam #13970) was used as the primary antibody with appropriate secondary antibody (anti-chicken AF488, 1:1000, Invitrogen #A11039) and DAPI Fluoromount-G® (Southern Biotech, #0100-20).

### Cortical neuron co-expression of GCaMP and mRuby

To assess co-expression of GCaMP and mRuby in cortical neurons, GCaMP expression was visualized via immunohistochemistry with anti-GFP, while mRuby’s intrinsic fluorescent properties were utilized for detection. Images were acquired using Olympus FV3000 confocal microscope with a 60x objective. In a single experimental animal, six distinct cortical regions of interest in layers II/III measuring 212 μm × 212 μm containing GCaMP and mRuby-positive neurons were selected and analyzed in ImageJ.

### Experimental design and statistical analysis

Statistics were performed using Prism 10 software (Graphpad) and R, and graphs were generated in Prism 10, R, and MATLAB. For all analyses, *n* refers to the number of animals, hippocampal slices, or neurons as is specified throughout. Normality tests were first performed to determine if data sets were normally distributed. For data sets that follow normal distribution, statistical significance was determined using either Student’s unpaired t-tests (indicated on graphs with asterisks, values defined in legends and results) or ANOVA (indicated by *p* values in legends as well as results). For multiple comparisons tests following ANOVA, multiplicity adjusted *p* values were reported (indicated on graphs with asterisks and defined in legends). Means and standard deviations were reported for all results unless otherwise specified. Survival analyses were completed to determine the hazard ratio and *p*-values as defined in the legends and results. Due to some animals being used for both control and OGD in vitro and animals from the same litter being used for sham and HI, a cox proportional hazard model was performed with clustering on mouse identifier.

### Data availability

All processed data associated with this study are present in the paper. All raw data is available upon reasonable request.

## RESULTS

### Injury Severity Optimization

Given the inherent challenges of chronic hippocampal imaging in vivo in an injured developing mouse pup, an in vitro preparation was utilized to track survival of hippocampal neurons after a mild insult. In vitro, previously published work from our lab tested multiple durations of OGD to identify a mild insult that triggered minor immediate neuronal death, manifest as quenching of fluorescence emission. This level of injury enabled longitudinal imaging of delayed neuronal death by tracking fluorescence emission in the surviving neurons (Bahari et al., 2024). With OGD durations ≥25 minutes, organotypic hippocampal slice cultures exhibited significant cell damage and early quenching. Neurons lost fluorescence emission after 2-3 hours of injury, prohibiting the study of chronic injury progression, whereas 10 minutes of OGD did not show evidence of any injury (no swelling, dying neurons, or caspase activation) (Bahari et al., 2024). 15-20 minutes of OGD was found to cause a small amount of caspase activation and neuronal swelling, but most neurons survived and continued to express fluorescent proteins (Bahari et al., 2024). Thus, this duration was selected for the experiments in this study to measure chronic injury after a mild insult in vitro (Fig. 1A). Similar to our previously published work, this duration of OGD resulted in a small amount of caspase activation (Fig. 2B) and a limited amount of immediate neuronal death (<10%, Fig. 3A). In vivo, a similar duration of hypoxia (15 min) after carotid ligation permitted longitudinal assessment of cortical neuron injury in the somatosensory cortex (Fig. 1, B and C). Longer durations of hypoxia resulted in higher mortality and more extensive acute quenching of neuronal fluorescence emission under the cranial window, prohibiting longitudinal imaging and quantification of injury progression with cellular resolution.

**Figure 3.**
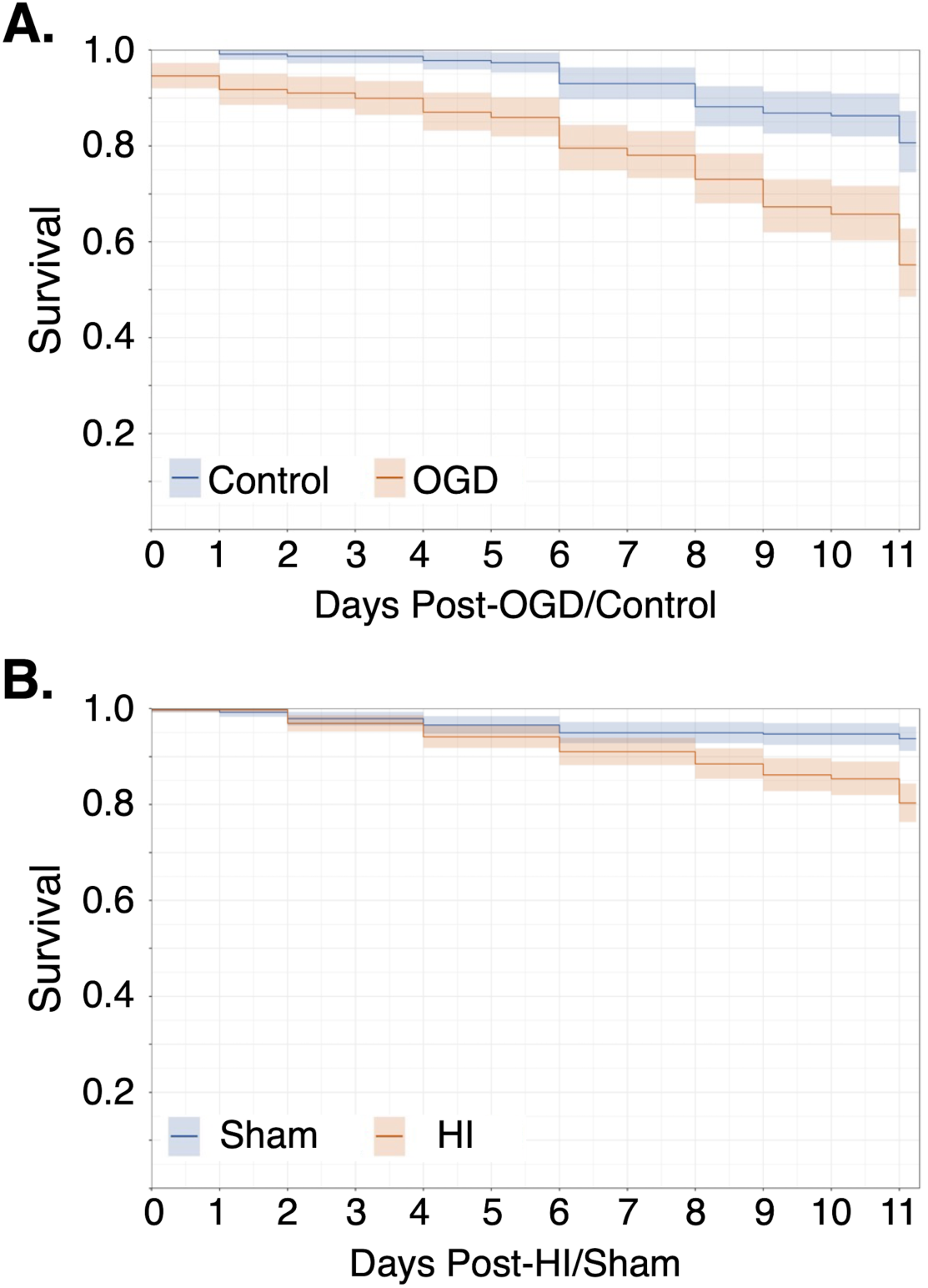
Persistent elevation of hippocampal and cortical neuronal death after mild perinatal hypoxia-ischemia. (A) In vitro hippocampal neuron Kaplan-Meier curve for control conditions (n=6 slices, 227 neurons, 26-49 neurons/slice) versus OGD (*n* = 6 slices, 277 neurons, 42-56 neurons/slice). The risk of neuronal death was significantly higher after OGD compared to control conditions (*p* < 0.001, hazard ratio 2.88 (95% CI = 1.6, 5.19)). (B) In vivo cortical neuron Kaplan-Meier curve for sham (*n* = 8 pups (5 female, 3 male), 376 neurons, 21-79 neurons/pup) vs. HI (*n* = 8 pups (3 female, 5 male), 389 neurons, 18-85 neurons/pup). The risk of neuronal death was significantly higher after mild HI compared to sham (*p* < 0.001, hazard ratio 3.242 (95% CI = 1.8, 5.84)).

### Neuronal death after mild HI was delayed until 2^nd^ week after injury

Using chronic 2P imaging of neurons expressing the constitutively fluorescent mRuby, hippocampal and cortical neuronal survival was manually tracked across z-planes in the same field of view using the fluorophore quenching assay after OGD in vitro (Fig. 2, A and B) and after HI in vivo (Fig. 2C and Supplemental Fig. 1). Ongoing emission of transgenically expressed fluorescent proteins has been shown to be a reliable biomarker of neuronal health in vitro (Arrasate et al., 2004; Balena et al., 2023), and can be tracked by longitudinal 2P imaging in vivo in the developing mouse cortex to identify neurons which undergo programmed neuronal death (Duan et al., 2020). Hippocampal pyramidal neurons that showed robust fluorophore expression after injury in vitro did not activate caspase activity acutely (Fig. 2B). When a neuron quenched in vitro, the caspase dye activated in the cytoplasm and then translocated to the nucleus where nuclear fragmentation was seen at 24h consistent with apoptotic neuronal death (Supplemental Fig. 2). The hazard ratio of death in vitro was 2.88 (95% CI = 1.6, 5.19, *p* < 0.001) after OGD and in vivo was 3.232 (95% CI = 1.8, 5.84, *p* < 0.001) after HI (Fig. 3, A and B). In both in vivo and in vitro experiments, the Kaplan-Meier curve shows that the progressive and persistently elevated neuronal death post-injury was most pronounced after the first week (Fig. 3, A and B). No sex differences were seen in the in vivo data; sex information was not available for the in vitro data to assess.

### No moderate or severe injury seen on ex-vivo magnetic resonance imaging

In vivo, in animals that survived 2 weeks post-HI or sham, no significant visible right hemispheric hypoplasia was seen by gross inspection (*n* = 8 HI (3 female, 5 male), 8 Sham (5 female, 3 male)). To confirm the absence of moderate to severe injury and validate this observation, ex vivo 14T, T2-weighted MRI in 3 HI (2 female, 1 male) and 3 sham animals (3 females) 18 days post-injury (P28) (Fig. 4A). There were no differences in the cortical, hippocampal, or cerebrum volumes and no volumetric asymmetries between the ipsilateral (right) and contralateral (left) cerebral hemispheres, cortices, or hippocampi in either sham or HI animals (Fig. 4B). In contrast, previously published cerebral and regional volumes measured with MRI after moderate to severe HI show significantly decreased volumes in the ipsilateral hemisphere compared to sham animals as early as 8 days post-injury (P18) (Burnsed et al., 2015). Our findings are consistent with both clinical data from patients with mild HIE, where MRI abnormalities, when present, are most often subtle (Prempunpong et al., 2018; Walsh and Inder, 2018), and with prior in vivo MRI studies in both preterm and in utero rodent models of mild perinatal insults showing no macroscopic injury 1-3 months after injury (Sanches et al., 2019; Cristancho et al., 2022).

**Figure 4.**
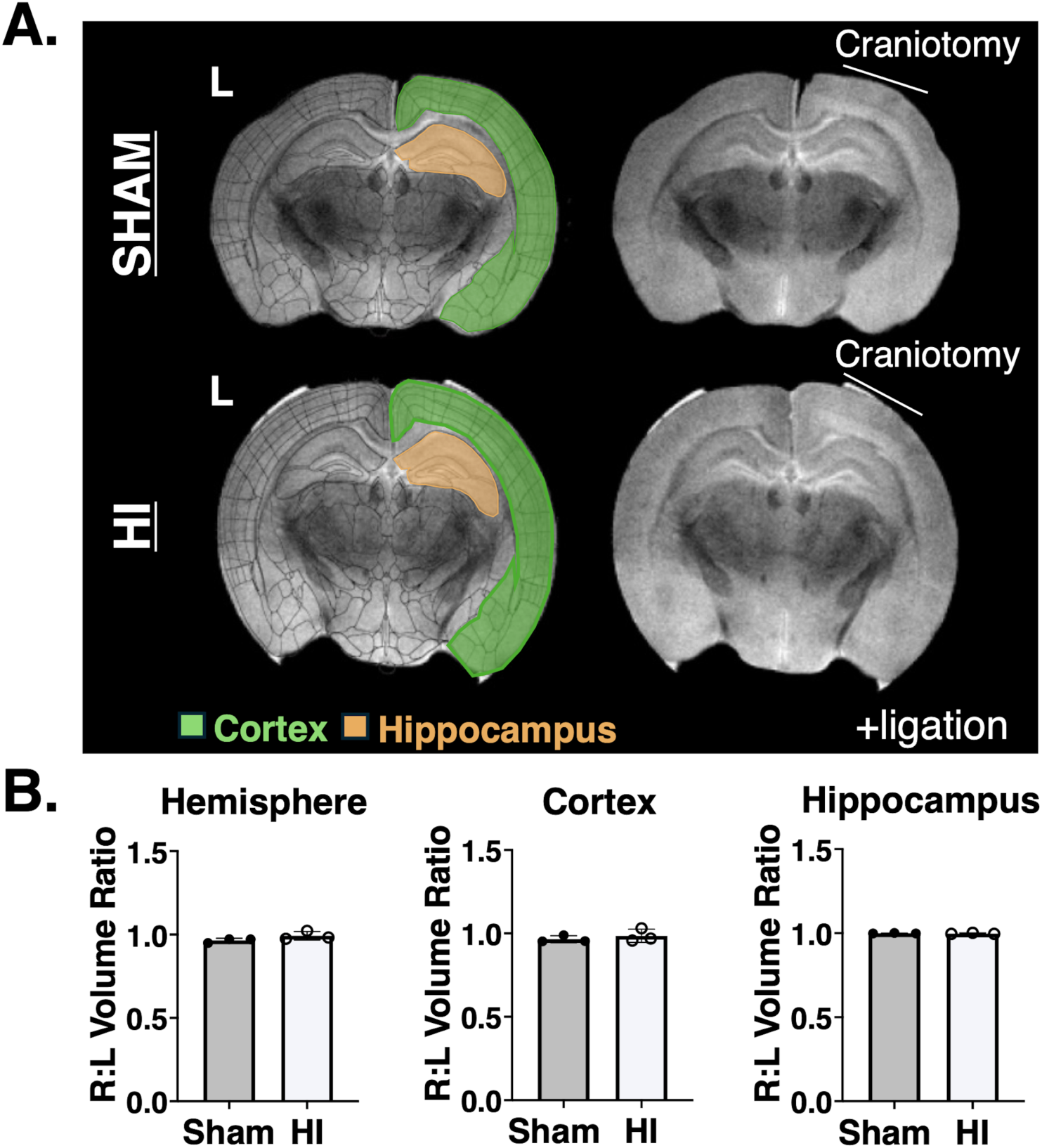
No moderate or severe injury detected by ex vivo MRI after mild perinatal hypoxic-ischemic injury. (A) Representative T2-weighted MRI coronal slices from sham and HI animals at location of 2P imaging. Overlays of the measured regions of interest for volumetric measures at that slice are shown. (B) No asymmetries were seen between cerebral hemispheres (*p* = 0.1586), cortices (*p* = 0.4778), or hippocampi (*p* = 0.5607) in sham (n=3, all female) or HI animals (*n* = 3, 2 female and 1 male) 18 days post-injury (P28; mean ± standard deviation, unpaired t-tests).

### Cortical network activity was transiently disrupted after mild hypoxic-ischemic injury

In human neonates, early cortical network activity is highly synchronous as seen in the well-known developmental EEG patterns of ‘trace discontinu’ and ‘trace alternant’ (Mizrahi and Hrachovy, 2016). Like humans, early murine cortical networks are highly synchronous with frequent spontaneous and responsive correlated network activity critical for normal maturation of cortical circuits (Khazipov et al., 2013; Huguenard et al., 2017). To measure the effects of mild HI on this cortical network activity in vivo, we performed unanesthetized head-fixed 2P neuronal GCaMP6s calcium imaging of layer II/III in the right somatosensory cortex through a cranial window. Consistent with clinical studies showing that up to 70% of neonates with mild HIE have abnormal EEG findings (Garvey et al., 2021), network activity was suppressed and decorrelated acutely and then recovered by 24 hours post-HI (*n* = 13 HI (7 female, 6 male), 13 Sham (7 female, 36 male), Fig. 5, A and B). At baseline, synchronous cortical network events were detected at a rate of 3.9 ± 1.2 events/minute in shams and 3.8 ± 0.5 events/minute after HI (Fig. 5B). The average duration of these events at baseline was 4.3 ± 1.6 seconds in shams and 4.2 ± 0.9 seconds after HI. The average proportion of co-active cells per network event, ‘Network Activation’, was 0.52 ± 0.09 in shams and 0.54 ± 0.12 after HI. One hour after HI, the frequency of, duration, and neuronal participation in network events decreased compared to baseline and sham, followed by full recovery at 24 hours after HI. No seizures were observed during the 2-hour imaging period after HI or during any later timepoints. No sex differences were seen in any of the metrics measured at any timepoint.

**Figure 5.**
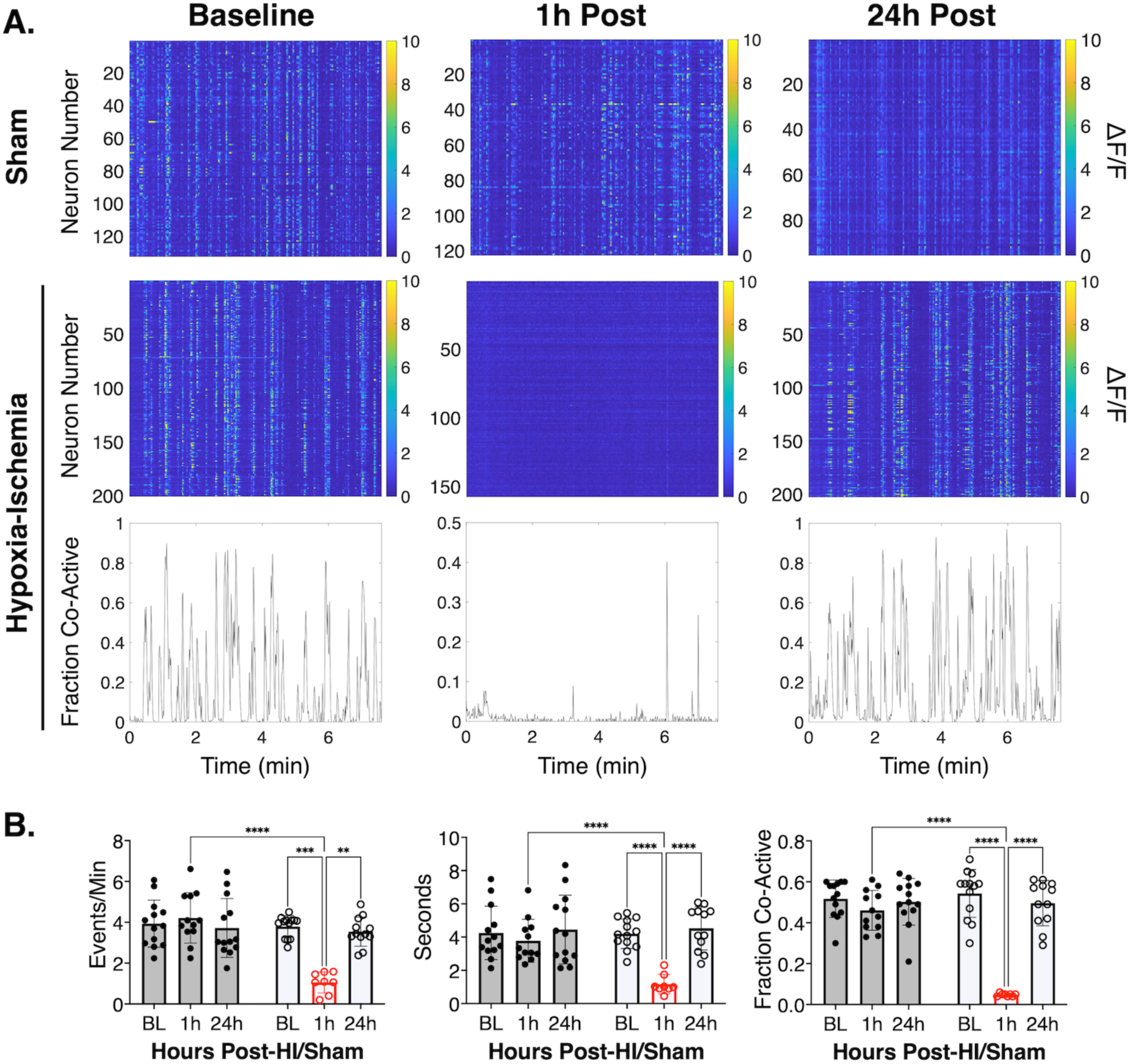
Mild perinatal hypoxia-ischemia transiently disrupted cortical network activity. (A) Representative raster plots of ΔF/F *syn*-GCaMP6s calcium signal taken at 1.1Hz from one sham and one HI animal at baseline, 1h post, and 24h post timepoints. Representative, corresponding traces to HI raster plots of the fraction of co-active neurons at baseline, 1h-, and 24h-post HI are also shown. (B) Mean ± standard deviation of network calcium activity metrics in HI (*n* = 13 (6 male, 7 female), open circles) versus sham (*n* = 13 (7 male, 6 female), filled circles) animals at baseline, 1h, and 24h timepoints. The frequency and duration of network events per animal is shown. Network activation is the average proportion of co-active cells per network event per animal. \*\*\*\**p* < 0.0001, *** *p* < 0.0001, ** *p* < 0.002, two-way ANOVA with post-hoc Tukey tests.

### Mild perinatal hypoxic-ischemic injury did not interfere with maturation of normal cortical activity

Longitudinal 2P calcium imaging in vivo showed that mild HI did not prevent maturation of normal cortical activity and functional assembly patterns 11 days after injury (Fig. 6). At baseline (P10) through 4 days (P14), the synchronous cortical activity shown in Figure 5 includes intrinsically generated network events which coexist with those triggered by sensory experience (including passive whisker deflections and spontaneous myoclonic twitches) (Khazipov et al., 2013; Che et al., 2018). By 11 days after injury (P21), cortical networks have decorrelated and the synchronous activity measured primarily represents functional neuronal assemblies that are triggered by receptive input such as active whisking and movement on the treadmill while the head-fixed animal was being imaged (Zagha et al., 2022). The frequency (*p* = 0.4782), duration (*p* = 0.7471), and proportional co-activation (*p* = 0.8123) of synchronous triggered cortical events were not different between sham and HI animals in vivo 11 days after injury (unpaired t-tests, Fig. 6). No sex-differences within or between the groups were seen.

**Figure 6.**
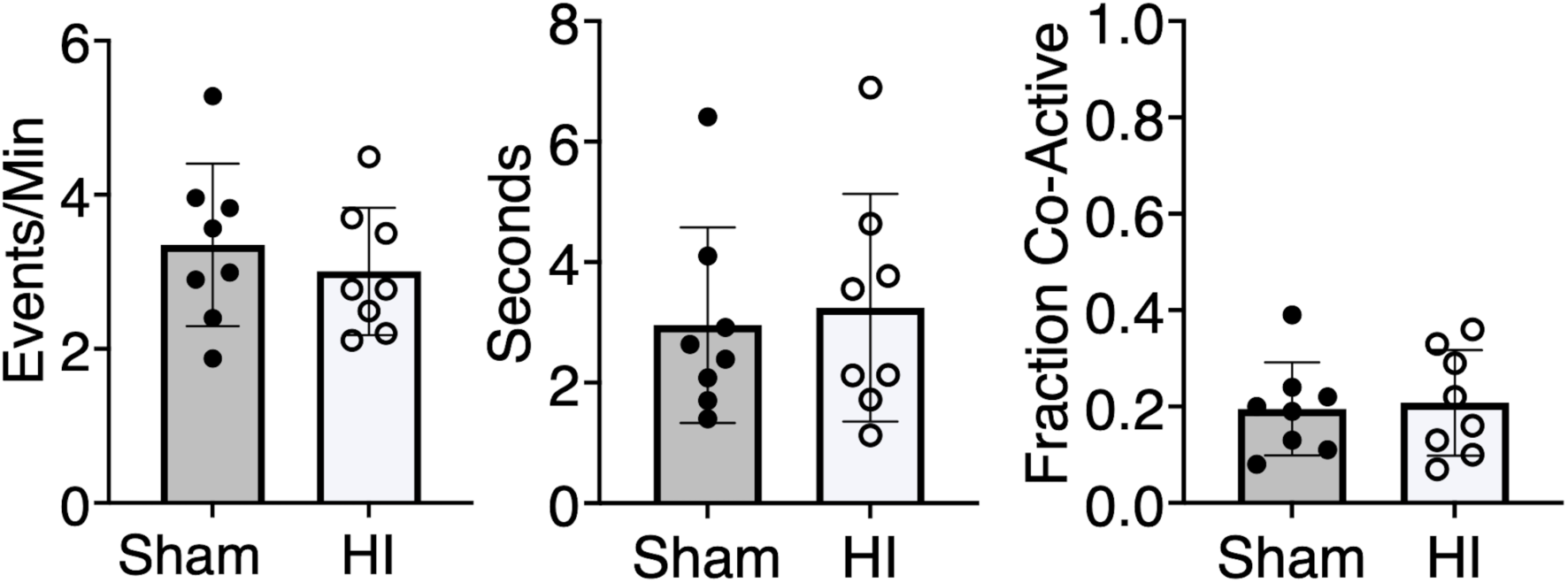
In vivo synchronous cortical activity was preserved 11 days after mild perinatal hypoxic-ischemic injury. Mean ± standard deviation of network calcium activity metrics in HI (*n* = 8 (3 female, 5 male), open circles) versus sham (*n* = 8 (5 female, 3 male), filled circles) animals 11 days after injury. The frequency (*p* = 0.4782) and duration (*p* = 0.7471) of cortical events per animal is shown. Cortical activation (*p* = 0.8123) represents the average proportion of co-active cells (‘Fraction Co-Active’) per event per animal. No differences were detected between groups (unpaired t-tests).

### Cortical neurons destined to die participated in physiological network activity for extended periods after injury

We hypothesized that cortical neurons destined to die 1-11 days after HI might undergo early synapse dysfunction and/or dendrite retraction leading up to their commitment to programmed cell death (Johansen et al., 1984; Yamamoto et al., 1986), thus interfering with their participation in synchronous network activity. To test this hypothesis in vivo, we measured the fraction of network events that each manually tracked neuron participated in (‘Neuronal Network Participation’) over time. The analyses were completed *by neuron*, controlling for correlation within pups. Repeated two-way ANOVAs were completed at *each* time point to measure effect of group and survival status. No significant differences were found at any timepoint based on survival status. At baseline (*p* = 0.369), 1- (*p* = 0.241), 2- (*p* = 0.962), and 4-days (*p* = 0.226) post-HI, network participation of *individual* neurons destined to die was not different from neurons that survived in sham and HI animals (two-way ANOVAs, Fig. 7). In neurons with ongoing fluorescent protein expression that died at later times after HI, their network activity participation was unaffected up to 4 days after mild injury. At 4 days (P14), the declining fractions of events in which neurons participate in after mild HI or sham reflects physiological desynchronization (Golshani et al., 2009; Che et al., 2018). Notably, mild HI did not disrupt this maturational process.

**Figure 7.**
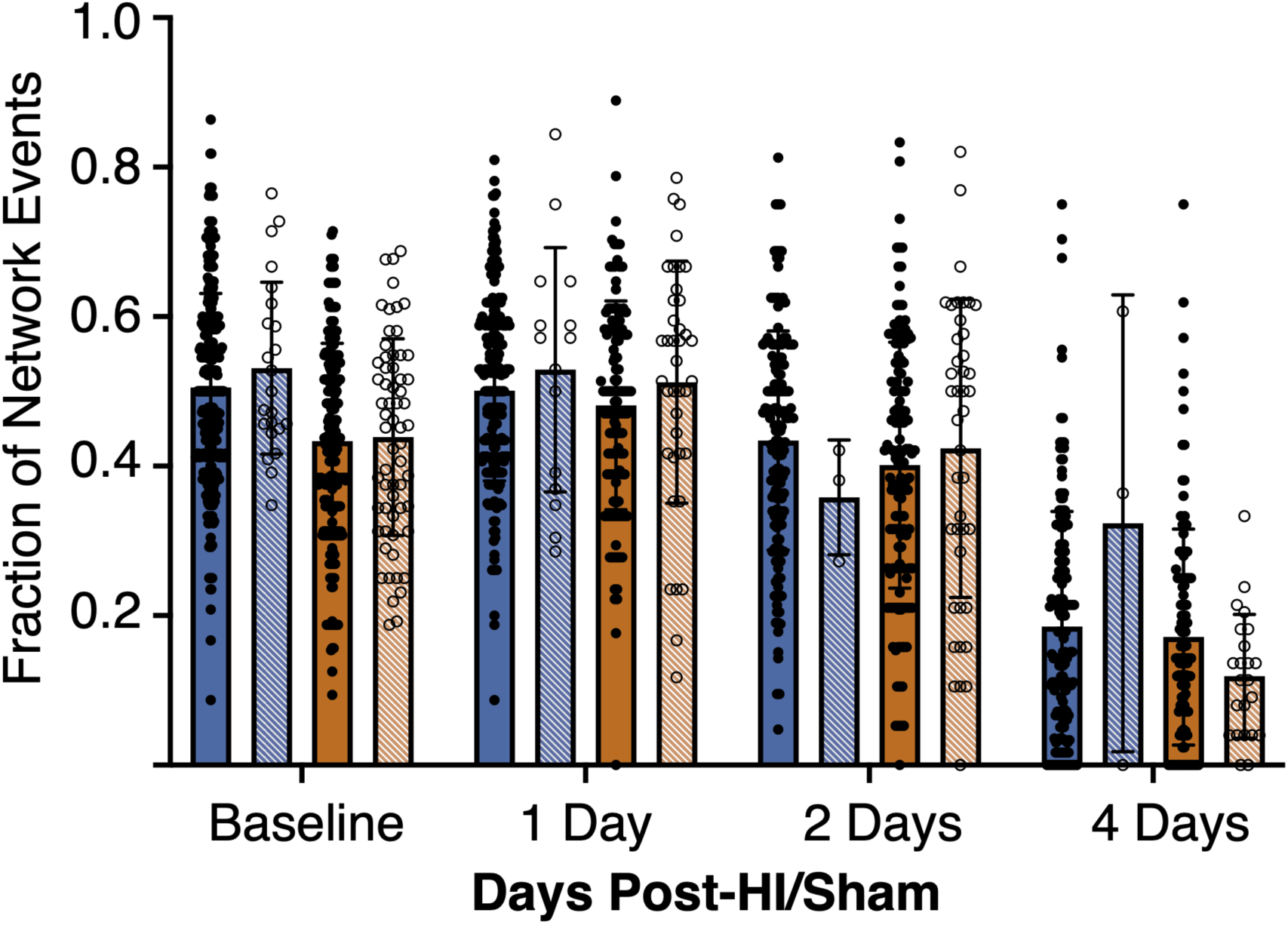
No difference in network participation of cortical neurons that die versus survive after mild perinatal hypoxia-ischemia in vivo. The fraction of total network events that *individual* neurons were active during (‘neuronal network participation’) was plotted for neurons that quenched prior to day 11 (‘died’, open circles) post-sham (blue striped) or post-HI (orange striped) versus those neurons that survive past day 11 (‘survived’, filled circles) post-sham (blue) or post-HI (orange). Data are shown as mean ± standard deviation at each timepoint after mild HI injury or sham for each group (*n* = 6-8 pups/group/timepoint, 3-333 neurons/group/timepoint), controlling for correlation within pups.

At baseline, there was a difference between mean neuronal network participation for the sham vs. HI group in the neurons that were tracked for survival analyses (*p* < 0.001, two-way ANOVA). As this may have been a confounding variable on survivability after HI or sham procedures, we completed a simple linear regression analysis on neuronal network participation using time as a predictor among the sham and HI groups. No relationship between network participation at baseline and time of neuronal cell death was detected (Supplemental Fig. 3). No differences in sham vs. HI group were found at later timepoints (*p* = 0.168 at 1 day, *p* = 0.108 at 2 days, and *p* = 0.234 at 4 days, two-way ANOVAs).

## DISCUSSION

In the present study, we showed that a mild perinatal HI injury, defined experimentally by the lack of widespread immediate neuronal death, caused a late and persistent elevation of hippocampal and cortical neuronal death through at least the second week after injury despite only transient disruption of cortical network activity. In vivo, for the first time, we showed full recovery of cortical network activity by 24 hours after mild HI. Mild HI did not affect maturational cortical network desynchronization or later cortical responses to sensory input. Additionally, mild HI did not result in frank volume loss on MRI 18 days after injury. Lastly, hippocampal and cortical neurons destined to die after mild HI showed markers of viability for up to 10 days after injury including robust fluorophore expression and full participation in network activity.

Early studies examining the impact of *mild* HI (5 minutes ischemia) on cortical and hippocampal neuronal activity in adult gerbils show 15-30 minutes of suppression followed by recovery with normal EEG activity detected for 48h (Suzuki et al., 1983). Contemporary studies of cortical EEG before, during, and for 2 hours after *moderate to severe* perinatal HI (carotid ligation and 45 minutes hypoxia) in neonatal mice show decreased EEG background amplitudes for 2 hours after injury (Burnsed et al., 2019). Notably, ∼50% of these animals develop seizures after moderate to severe HI. Suppression of network activity in the 2 hours after mild HI with subsequent recovery by 24 hours (Fig. 3) is consistent with both this early and contemporary work. We validated these findings with cellular resolution imaging of cortical networks to measure the recovery of this activity after mild HI for the first time. No acute seizures were seen. This lack of seizures is a critical defining feature of human mild HIE.

The possibility that neurons survive initial insults only to succumb later is of great interest because it suggests an extended therapeutic window for neuroprotective interventions. In early studies, ‘delayed’ neuronal death after ischemia in rodents is seen maximally 48-72 hours after injury (Kirino, 1982; Pulsinelli et al., 1982). At a gross level in a perinatal injury model employing prolonged hypoxia and ischemia, there is ongoing, progressive cerebral volume loss for months, raising the possibility of ongoing neuronal death (Geddes et al., 2001). However, efferocytosis of dying neurons and scar formation and retraction likely also contribute to these changes. Subsequent studies have demonstrated dying neurons histopathologically for up to 7 days after moderate to severe HI (Nakajima et al., 2000; Northington et al., 2001, 2011b). However, these single time-point histological assays cannot determine the precise time point at which neurons fully commit to cell death and are beyond rescue (Zille et al., 2012; Balena et al., 2023). This is the first definitive demonstration of ongoing, new, irreversible commitment to programmed neuronal death up to 11 days after mild HI. The lack of volumetric changes on MRI 18 days after injury confirmed this was a mild injury, as mice exposed to longer periods of hypoxia have well-established volume changes in the ipsilateral hemisphere detected by MRI by this age (Burnsed et al., 2015). Additionally, the absence of macroscopic abnormalities at the timepoint studied does not rule out loss of tissue volumes at later times. It is remarkable that the fluorophore quenching assay was sensitive enough to detect the mild cortical injury at this time. Longer-term MRI studies are needed to determine the impact of mild HI on brain development and organization, including functional connectivity. Persistent elevation of neuronal death for weeks or months after injury could lead to increased network simplification which may be pathological and contribute to the long-term cognitive deficits seen at later ages in these patients.

Studying neuronal death after injury using histopathological assays is complex given dynamic changes in the number of neurons that are “visibly dying” on these assays, the rate of commitment to programmed cell death prior to the assay, the rate of efferocytosis of dying neurons, and the time elapsed since injury (Balena et al., 2023). Tracking individual neurons in real time over days and weeks allowed us to overcome a number of these limitations (Linsley et al., 2019). Progressive injury has been hypothesized for decades after perinatal HI, and this progression forms the theoretical basis for the evolving depth of encephalopathy seen clinically after injury in both perinatal (Sarnat and Sarnat, 1976) and mature (Levy et al., 1985) brains. The current study evaluated only mild HI injury, but it was the first to definitively show that progression on a cellular level with a prolonged time window. Perhaps the most important new finding is that the neurons that died late after HI showed evidence of intact function and strong viability on both the single cell and network level. These neurons participated in physiological network activity and demonstrated normal maturation for at least 4 days after injury. These neurons demonstrated functional translational machinery to produce transgenic proteins, absence of caspase activation acutely, and functional connections to the cortical network indicating robust ongoing synaptic activity. In total, the findings suggest an extended therapeutic window for mild HI, as these neurons were likely salvageable for many days after injury.

The cellular mechanisms of late neuronal death after mild HI remain to be discovered. Developmental neuronal apoptosis is largely complete at the time points studied (Southwell et al., 2012; Kuan Wong and Marín, 2019), although a recrudescence of these processes is a possible driver. It is also possible that mild HI exacerbated isoflurane related neurotoxicity in our animals. Prior studies of P7 mouse pups exposed to 5 hours of isoflurane demonstrated significant apoptotic neuronal death in cortical neurons immediately after exposure (Istaphanous et al., 2013). Our HI and sham pups were equally exposed to <1 hour of isoflurane at P7-8, and then another 5-10 minutes at P10. This is a lower overall exposure than used by Istaphanous et al.(2013), but this remains a possible confounding variable. Another consideration is that our cellular-level recordings did not suggest an activity-dependent mechanism of cell death (Wong et al., 2018; Duan et al., 2020). The mechanisms of programmed neuronal cell death in the developing brain are diverse and the subject of intense study (Northington et al., 2011a; Pfisterer and Khodosevich, 2017; Davidson et al., 2021). Future studies will need to determine how crucial these various programs are to late neuronal death after mild HI. Understanding these mechanisms is critical for evaluation of therapeutic options.

Our observations regarding late neuronal death were subject to practical limitations of the duration and locations where neuronal death and activity could be tracked in the developing brain. We did not detect any detrimental effects of the persistent elevation of neuronal death on cortical activity 11 days after injury, or macroscopically by MRI at 18 days after injury, that might explain the later neurodevelopmental deficits seen in human neonates after mild HIE (Conway et al., 2018). However, the metrics of network activity we measured may have missed important developmental transitions that will require more detailed analyses. Additionally, the present study lacks long-term behavioral outcomes. Future studies can test whether neurobehavioral deficits are present in these animals. Notably, two rodent models of mild perinatal injury (one preterm mild HI (Sanches et al., 2019) and one in-utero mild hypoxia (Cristancho et al., 2022)), which showed no abnormalities on MRI 1-3 months after injury, *did* demonstrate long term behavioral deficits. After our mild injury in vivo, it is possible that MRI with advanced sequencing at later timepoints may be more sensitive for any long-term anatomical consequences of mild HI, and subsequent behavioral testing may find correlates to the human deficits seen. The ‘normal’ cortical activity we measured at 11 days after injury does not predict whether later dysmaturation occurs. Additionally, in vivo we assessed the somatosensory cortex rather than the hippocampus, which is the most vulnerable region after moderate to severe HI (Burnsed et al., 2015) and we may have missed more abundant and different effects on cell death progression and network abnormalities occurring there.

Despite these limitations, our data provide a strong rationale for future studies looking at later timepoints and long-term behavioral outcomes. Additionally, we provide an in vivo benchmark for how long the therapeutic window is open after mild HI and against which therapeutic interventions can be tested to see whether they attenuate ongoing neuronal death. Using the methods developed in this study, one can also determine whether there are functional metrics that predict persistent neuronal death and/or biomarkers of the severity of persistent neuronal death. Since there is likely similar long-term progressive neuronal death occurring in our neonates with mild HIE, acute MRI while the patients are still in the NICU may miss the extent of an injury and not avail them to developing therapeutic interventions. Later repeat neuroimaging is the minimum that can be done to begin to ascertain the actual neurodevelopmental burden of mild HIE. Importantly, this evidence for late, sustained neuronal death following mild HI must be incorporated in decisions regarding possible therapeutic interventions in these patients.

## ACKNOWLEDGEMENTS

We acknowledge our funding sources: National Institute of Health grant K08 NS121599-01 (M.A.M) and grant R35 NS116852 (K.J.S).

We acknowledge and thank Dr. A. Cruz-Martin for his generous time training in the cranial window placement on murine pups.

