## Supplemental Figures for "Ongoing loss of viable neurons for weeks after mild perinatal hypoxia-ischemia"

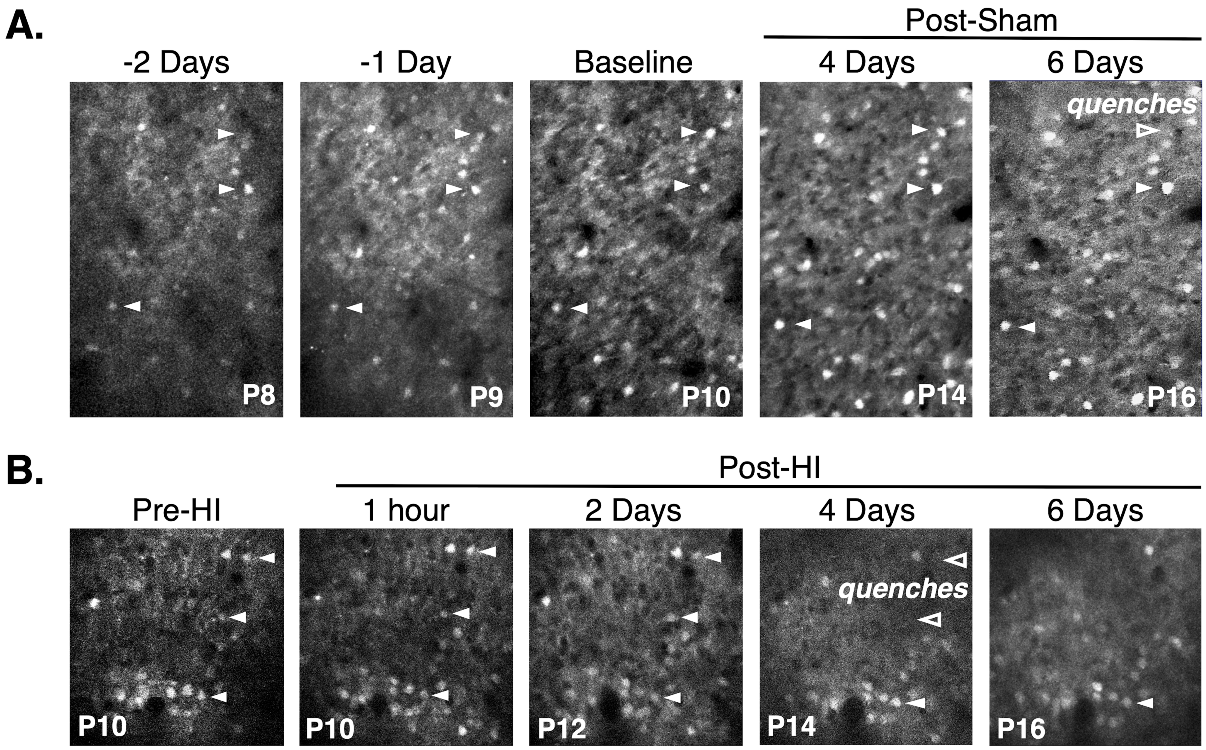


**Supplemental Figure 1 Longitudinal tracking of cortical neuron survival in vivo after sham and HI.** Unanesthetized, in vivo 2P fluorescence images of layer II/III somatosensory cortex over time demonstrating tracking of mRuby-positive cortical neurons from a sham (A) and HI (B) animal. White arrowheads show representative tracked neurons. Open white arrowheads show area where tracked neuron has quenched.

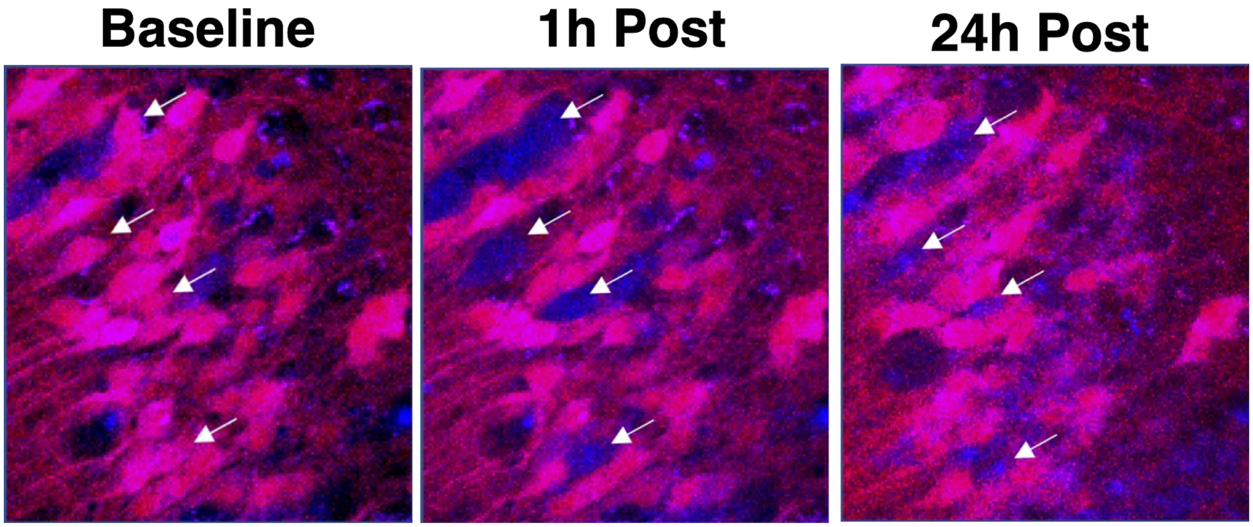


**Supplemental Figure 2 Caspase activation in dying hippocampal pyramidal neurons.** Representative 2P fluorescence images of mRuby-positive hippocampal pyramidal neurons (pink) in vitro incubated with NucView® Blue Caspase-3 dye (blue) in an unhealthy DIV16 organotypic hippocampal slice. Neurons that quenched in unhealthy conditions (arrows) show initial diffuse, cytoplasmic caspase staining and finally nuclear localization showing fragmented nuclear material consistent with apoptotic cell death.

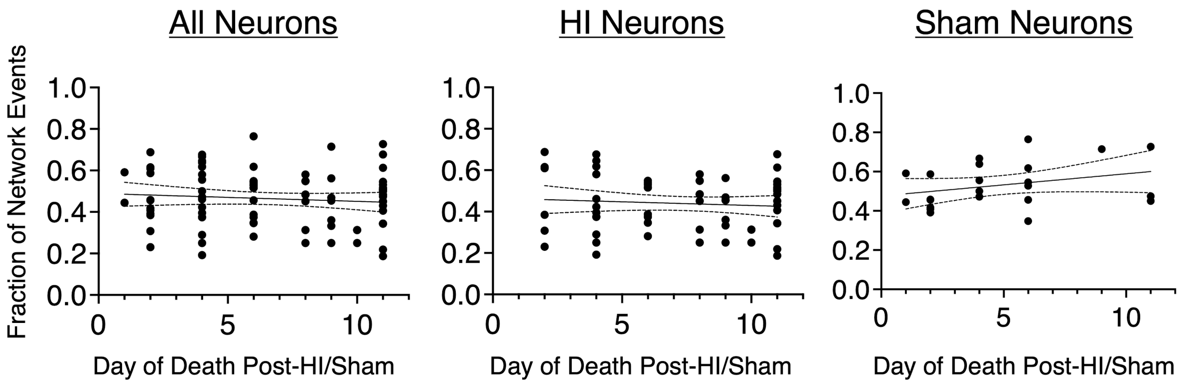


**Supplemental Figure 3 Baseline neuronal network participation was not predictive of later neuronal death.** Simple linear regressions of neuronal network participation at baseline (P10) vs. day of death are shown for all neurons (16 pups, 85 neurons; $R^2 = 0.01$, $p = 0.39$), HI neurons (8 pups, 62 neurons; $R^2 = 0.01$, $p = 0.51$), and sham neurons (8 pups, 23 neurons; $R^2 = 0.09$, $p = 0.15$). 95$^{th}$ percentile confidence limits shown with dashed lines.
